# Red-wine gene networks associated with exceptional longevity in humans

**DOI:** 10.1101/2025.06.21.660865

**Authors:** Patricia Lacayo, Alexandria Martignoni, Kenneth Park, Christianne Castro, Shin Murakami

**Affiliations:** Department of Foundational Biomedical Sciences, College of Osteopathic Medicine, Touro University California, Vallejo, CA, USA

**Keywords:** aging, longevity, centenarian, alternative medicine, polymorphism, red wine, polyphenol, resveratrol, catequins, rotundone, garlic acids

## Abstract

Moderate consumption of red wine has been associated with healthy aging and longevity, defined as one drink per day for women and two drinks per day for men (approximately 142 ml or 5 oz per drink). Previous research has revealed the health benefits of red wine, particularly in relation to cardiovascular disease. However, the influence of genetic factors on these benefits remains to be elucidated. In this study, we explored genes linked to red wine and created a curated gene set that intersects with those related to centenarians, who are markers of exceptional longevity. Utilizing over 190 databases, we identified and validated a curated list of genes, and conducted gene set enrichment analysis as well as enrichment analysis of annotations and diseases. Our findings highlighted 43 genes connected to centenarians, suggesting that these genes play a crucial role in stress response and apoptosis, which are essential for cell survival and renewal. Additionally, we noted that these genes were enriched in pathways associated with smooth muscle cell proliferation, neuroinflammation, nucleotide excision repair, and lipoprotein metabolism (false discovery rate, FDR < 3 × 10^−07^). Gene set enrichment analysis indicated significant tissue expression in the gastrointestinal, cardiovascular, and respiratory systems. Furthermore, the disease-gene enrichment analysis pointed to associations with the diseases related to the tissues, including cardiovascular disease (heart disease and stroke), type 2 diabetes, gastrointestinal diseases and metabolic diseases, immune diseases, and cancer (FDR < 9.37 × 10^−6^); notably, cardiovascular diseases, diabetes, and cancer are leading causes of death, suggesting that the genes may be protective against those diseases. Although further research is necessary to uncover additional genes, this study provides the first genetic overview of the health benefits of red wine, emphasizing its potential in supporting healthy aging and longevity.

## Introduction

The field of biological aging has made remarkable progress in uncovering the cellular and molecular mechanisms underlying age-related diseases (Brunet, 2020; Sziraki et al., 2023; Balmorez et al., 2023). Research on centenarian populations offers a unique perspective, guiding scientists in their efforts to identify factors that may promote longevity. However, a significant challenge persists: the relatively limited exploration of why centenarians demonstrate exceptional resistance to age-related diseases. A particularly intriguing phenomenon known as the French paradox (Renaud and de Lorgeril, 1992; Ferrières et al., 2004) has garnered substantial scientific interest, particularly in light of the exceptional longevity exhibited by Jeanne Calment, a French centenarian who lived to the age of 122 years (Robine et al., 2019; Robin-Champigneul et al., 2020). Epidemiological studies have consistently demonstrated that the French population has a lower incidence of coronary heart disease (CHD), a leading cause of death, despite a dietary pattern characterized by a higher intake of saturated fats (Ferrières et al., 2004). This observation raises questions regarding the protective factors that may be implicated, with a notable emphasis on red wine consumption.

Recent studies have increasingly explored the potential health benefits of red wine. Moderate consumption of red wine has been associated with reduced risks of cardiovascular disease, cancer, and improved cognitive function; moderate consumption is defined as one drink per day for women and two drinks per day for men (142 ml or 5 oz per drink) (NIAAA, 2023; US dietary guidelines, 2023; Wojtowicz et al., 2023). Red wine has been found to reduce atherosclerosis-related inflammatory markers in healthy individuals (Sheng et al., 2024), inhibit aromatase to potentially benefit patients with breast cancer (Pergolizzi et al., 2024), and improve cognitive function when included in the diet (Boushey et al., 2020). Furthermore, red wine has been shown to reduce markers of oxidative stress, inflammation, and nephropathy (Lombardo et al., 2023). Although previous studies have established the potential health benefits of red wine, the relationship between red wine and genes remains to be explored.

Red wine contains a wide variety of biomolecules, such as catequins, rotundone, garlic acids, resveratrol (3,5,4’-trihydroxystilbene), and other polyphenols (Markoski et al., 2016), which are known to interact with genes and gene products. Cagtequins are scavengers of reactive oxygen species and can inhibit pro-oxidant enzymes (Bernatoniene J, Kopustinskiene, 2018).

Resveratrol, a well-characterized component of red wine, targets SIRT1, a sirtuinof the Sirtuin family that encodes nicotinamide adenine dinucleotide-dependent deacetylase. Although SIRT1 function has yet to be identified in humans, sirtuin proteins in yeast regulate epigenetic gene silencing and suppress rDNA recombination (Wu et al., 2022). SIRT1 is associated with reduced risks of hypertension, diabetes (i.e., reducing hemoglobin A1C), protective vascular function, metabolic syndrome, and cardiomyopathy (Hausenblas et al. 2015; Kane and Sinclair, 2018).

Resveratrol increases the level of Sirtuin-1 by inhibiting the TLR4/NF-κB/STAT signal cascade and increases the level of brain-derived neurotrophic factor (BDNF) in the blood NOS-3 (nitric oxide synthase-3) activity (Wiciński et al., 2018). Red wine polyphenols suppress the secretion of ApoB (ApoB48 and ApoB100), affecting lipid metabolism (Pal et al., 2003 and 2005).

Although mounting evidence suggests the health benefits of red wine, a red-wine component, resveratrol, may have no effects on cardiovascular health in clinical trials where the beneficial dose of resveratrol is much higher than the dose found in red wine (Bonnefont-Rousselot 2016).

A comprehensive genetic analysis of the health benefits associated with red wine is expected to enhance our understanding of its positive effects on health. In this study, we systematically investigated the genes that are upregulated or downregulated following the consumption of red wine and its components. Among the red wine genes, we identified genes associated with centenarians, serving as indicators of exceptional longevity. Our research aims to address existing knowledge gaps by compiling a gene set pertinent to red wine, delineating genetic networks linked to moderate red wine consumption, and elucidating the implications of these findings for human longevity and health benefits.

## Materials and Methods

### Study Design and Gene List Generation

This study was designed to identify and analyze genes associated with longevity by examining the overlap between the genetic profiles of red wine consumers and centenarians. Gene identification was performed as previously described (Machino et al., 2014; Vahdati et al., 2017; Balmorez et al., 2023; Baidal et al., 2024). Briefly, a comprehensive gene list was created using the keywords “red wine” and “centenarians” in the GeneCards database (www.genecards.org; last accessed 3/18/2025), which aggregates data from over 190 databases and references relevant scientific literature concerning human gene entries. The genes were ranked according to the Boolean model (Tenopir, 2008) to identify the corresponding documents, and a method known as the practical scoring function was used to assess relevance. The retrieved data formed a primary gene list for further validation.

### Gene Review and Validation

Two independent investigators reviewed the literature corresponding to each gene identified, and the consensus of the three reviews was obtained. The inclusion criteria were as follows: (1) the literature was accessible; (2) each gene had been the focus of research in the cited studies; and (3) the result was statistically significant (p < 0.05). This ensured that only relevant studies were considered for analysis. The genes that exhibited inconsistencies across these reviews underwent a comprehensive validation process and were included only when the review demonstrated statistically significant outcomes (p < 0.05). Studies lacking confidence levels of p-values and/or any other statistical criteria, as well as those with inaccessible related research were excluded from the analysis.

### AI Utilization for Literature Summarization

The use of a generative AI chatbot (Version ChatGPT-4o; https://chatgpt.org/; last accessed on 1/05/2025) was tested to evaluate its capacity to streamline the research. The information of each study manuscript, including the abstract, methods, and results, was used to generate summaries. These summaries were then reviewed, compared with the manual reviews, and validated for consistency with the study’s objectives. It is important to note that this process was conducted separately from the primary research and did not influence the study outcomes.

### Data Analysis

We used multi-omics analyses with the gene set generated in this study. Gene set enrichment analysis (GSEA) was performed using STRING 12.5 (Szklarczyk et al., 2025) (https://string-db.org/, last accessed on 3/8/2025) as described by (Balmorez et al. (2023) and Baidal et al. (2024). Network topology and over-representation (ORA) analyses were performed using the web-based gene set analysis toolkit, WebGestalt 2024 (Elizarraras et al., 2024) (https://www.webgestalt.org/, last accessed on 3/8/2025). Network Topology Analysis examines the characteristics of the identified genes, while ORA assesses the presence of genes in various categories, including biological, cellular, and molecular functions, as well as pathways (such as Reactome), tissues, and disease categories. The network database used was PPI BIOGRID (Elizarraras et al., 2024). We used statistical significance, as described in the text.

## Results

Using approximately 180 databases (Komal et al.,2024), a total of 245 genes were found to be associated with the keyword “Red Wine,” while 627 genes were associated with “Centenarian.” This analysis yielded 72 genes that were concurrently linked to both “Red Wine” and “Centenarian” (Materials and Methods) (Figure 1).

**Figure 1.**
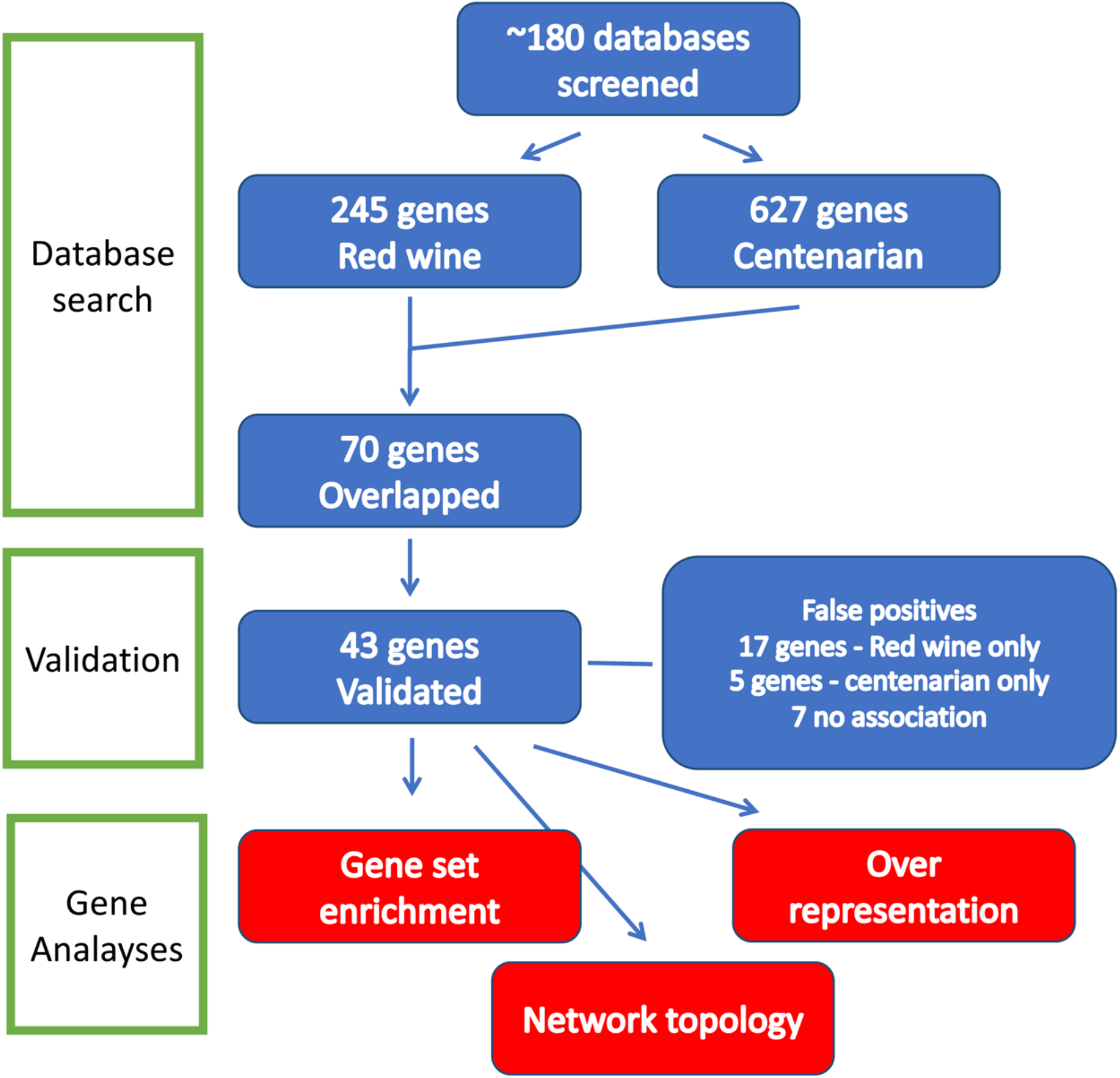
Flow diagram of gene identification and analysis. Three types of analyses, including network topology analysis, over-representation analysis (ORA), and gene set enrichment analysis (GSGA), were performed. See text for details.

To confirm these findings, 72 genes were validated through three independent reviews. In cases where evidence was insufficient, such as instances of both upregulation and downregulation, a conservative statistical threshold of p < 0.05 was applied to establish significance. The validation process identified genes that did not meet the validation criteria and were categorized as false positives: five genes (6.9%) associated with red wine, seventeen genes (23.6%) associated with centenarians, and seven genes (9.7%) linked to both categories. We also validated using the AI chatbot, which resulted in 48 genes, including five false positives (10% error rate). In summary, we confirmed the existence of 43 genes associated with both red wine and centenarians, of which 40 were recognized by the analytical programs employed in this study (Materials and Methods).

We conducted three types of analyses, as illustrated in Figure 1: network topology analysis, over-representation analysis (ORA), and gene set enrichment analysis (GSEA). Both ORA and GSEA evaluate the likelihood of overrepresentation within specific biological pathways. ORA calculates the number of genes from a given pathway included in a particular list, whereas GSEA analyzes the distribution of these pathway genes within a ranked list.

### Network topology analysis and over-representation analysis

Utilizing a threshold of false discovery rate (FDR) < 10 × 10^−5^, we conducted multiple analyses of the gene set, as shown in Figure 2. Figures 2A and 2B present the outcomes of the network topology analysis, while Figure 2C depicts gene over-representation by quantity. The analysis encompassed a search against 2,194 genes within the networks, incorporating the top 10 neighboring genes (Figure 2A). The findings indicate an intricate interconnection among the analyzed genes, but not diverse genes unrelated to each other.

**Figure 2.**
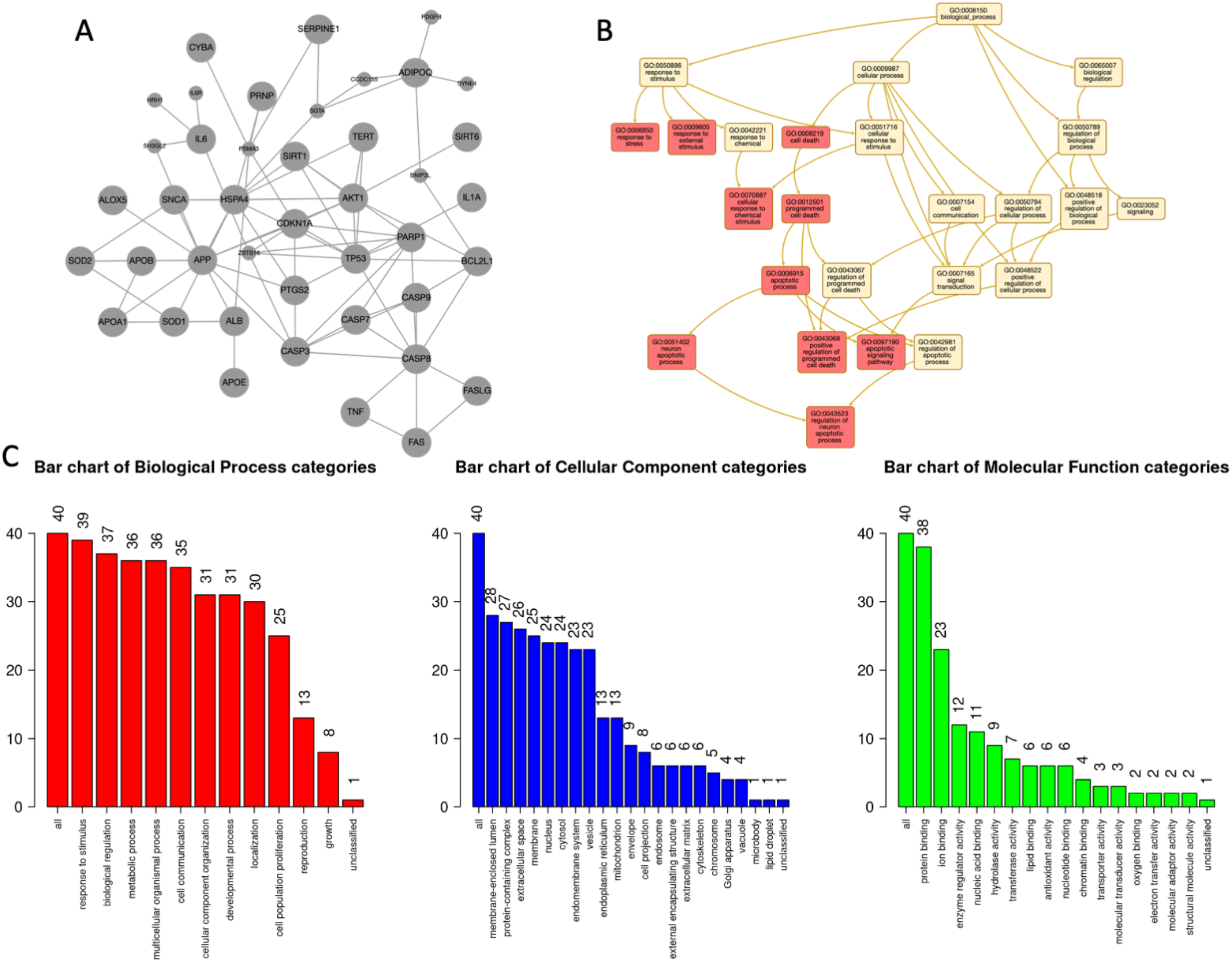
Nature of the genes associated with red wine and centenarians. A. Network topology analysis of gene networks (large gray nodes, seed genes; small gray nodes, top-ranking neighbors). The program (Webgestalt) generated minor variations in the figure despite the same statistical results. An example is shown. B. Network topology analysis of enriched GO terms (orange-pink, enriched GO terms; yellow, ancestors of the terms). C. Over-presentation analysis using gene ontology and the processes shown in the figure (red, biological category; cellular, cellular category; and green, molecular function categories). The numbers of each category are shown, and those with over 25 genes are listed in the text. Larger figures have been included (Link to be assigned by the journal).

**Figure 3.**
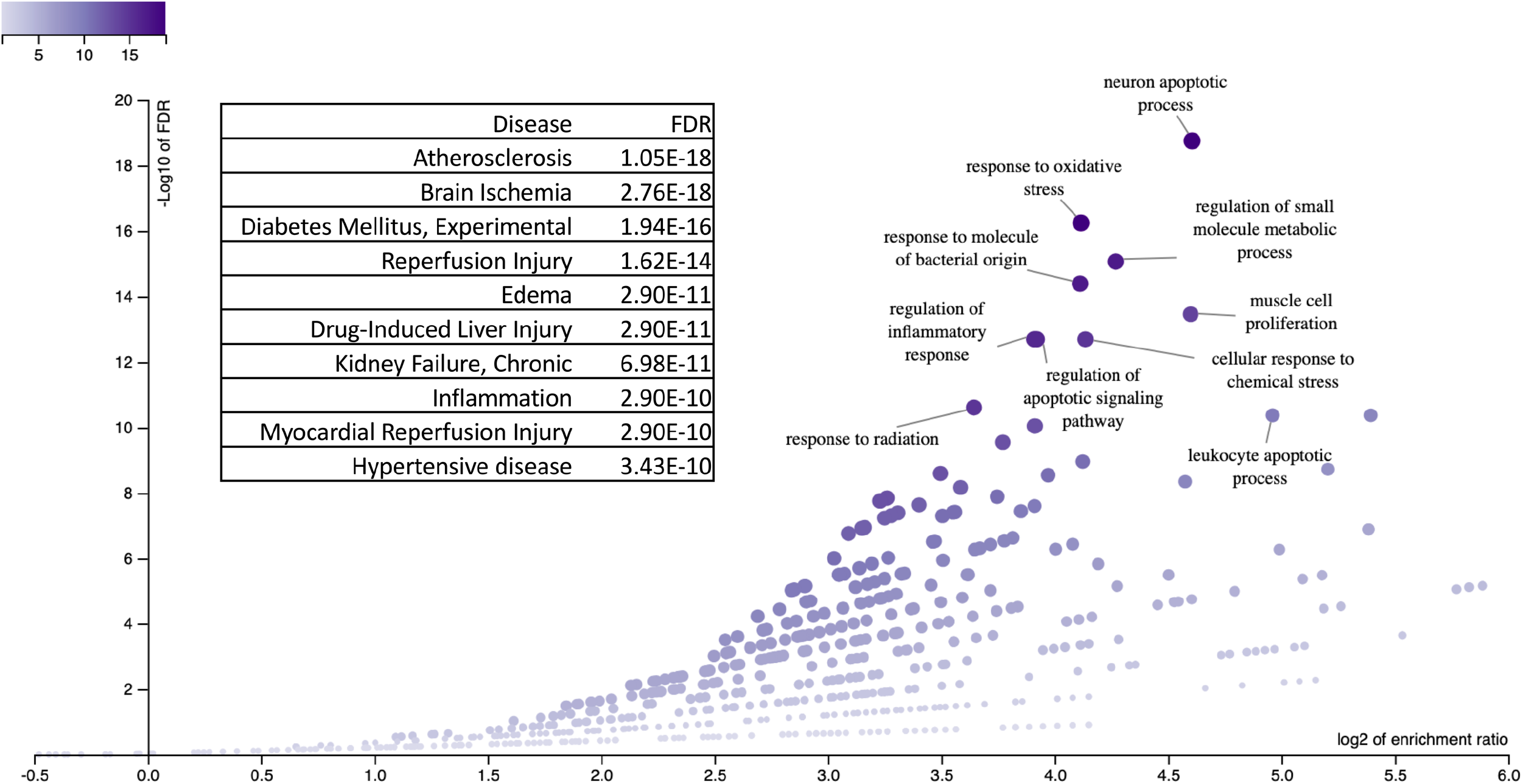
Over-representation analysis of disease categories. The top 10 disease categories are labeled and shown in the box with the statistics. Over-representation analysis was performed.

Figure 2B summarizes the enriched top 10 Gene Ontology (GO) categories, which are classified into two primary groups: (1) response to stress and stimuli, with an adjusted p-value < 4.2 × 10^−13^, and (2) apoptosis/cell death, with an adjusted p-value < 6.1 × 10^−13^. Figure 2C shows the gene distribution across three domains: (1) biological processes, comprising eight categories with over 25 genes per category, summarized as follows: response to stimulus (39 genes), biological functions (37 genes), metabolic processes (36 genes), cell communication (35 genes), cellular component organization (31 genes), and localization (30 genes); (2) cellular components, which includes four categories with over 25 genes per group, such as membrane-enriched lumen (28 genes), protein-containing complex (27 genes), extracellular space (26 genes), and membrane (25 genes); and (3) molecular function, characterized by a single category comprising over 25 genes, specifically protein binding (38 genes).

The over-representation analysis of the disease enrichment showed a significant enrichment with leading causes of death (Figure 4), including cardiovascular problems (arteriosclerosis, brain ischemia, reperfusion injury, hypertension, and other artery diseases), diabetes relevant to experimental systems, immune problems and inflammation, and kidney failure (FDR < 3.4 × 10^−10^). The list of diseases largely overlapped with the disease-enrichment results obtained by another method, gene set enrichment analysis (below).

**Figure 4.**
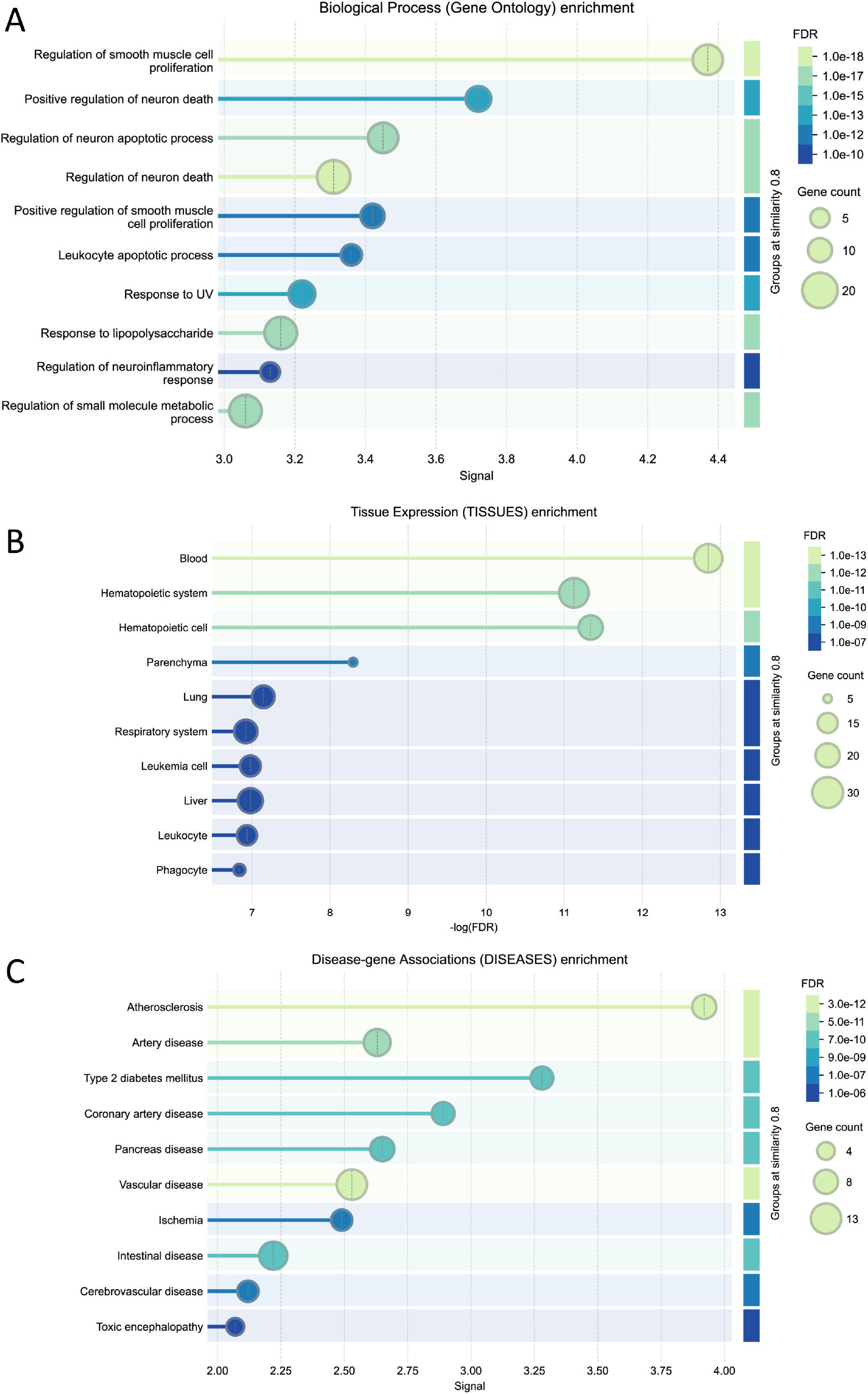
Gene set enrichment analysis (GSEA) A. Biological ontology pathway. B. Tissue expression. C. Disease-gene association classification. STRING-DB was used to visualize the grouped and compiled categories of the identified proteins. The top 10 categories were grouped using a similarity score of ≥ 0.8. Supplementary material is available for a complete list of categories.

### Gene set enrichment analysis

To further validate the results above, we used gene set enrichment analysis to investigate biological pathways, tissue expression, and disease-gene associations as follows.

#### Biological pathways

A total of 324 biological ontology pathways were identified (Supplementary Table 1), with redundancies eliminated based on their similarity (refer to Materials and Methods). Figure 4A presents a summary of the top 10 identified pathways, which include: (1) smooth muscle cell proliferation, recognized for its diverse roles across the gastrointestinal, cardiovascular, renal, and respiratory systems; (2) neuron-related pathways encompassing processes such as neural death and neuroinflammatory responses; (3) leukocyte apoptosis; (4) responses to ultraviolet light, specifically those involving nucleotide damage response and excision repair mechanisms; and (5) metabolic processes associated with lipoproteins. Notably, gene set enrichment analysis provided a more comprehensive understanding of the pathways compared to over-representation analysis, which primarily highlighted responses to stress and stimuli, as well as apoptosis and cell death.

**Table 1.**
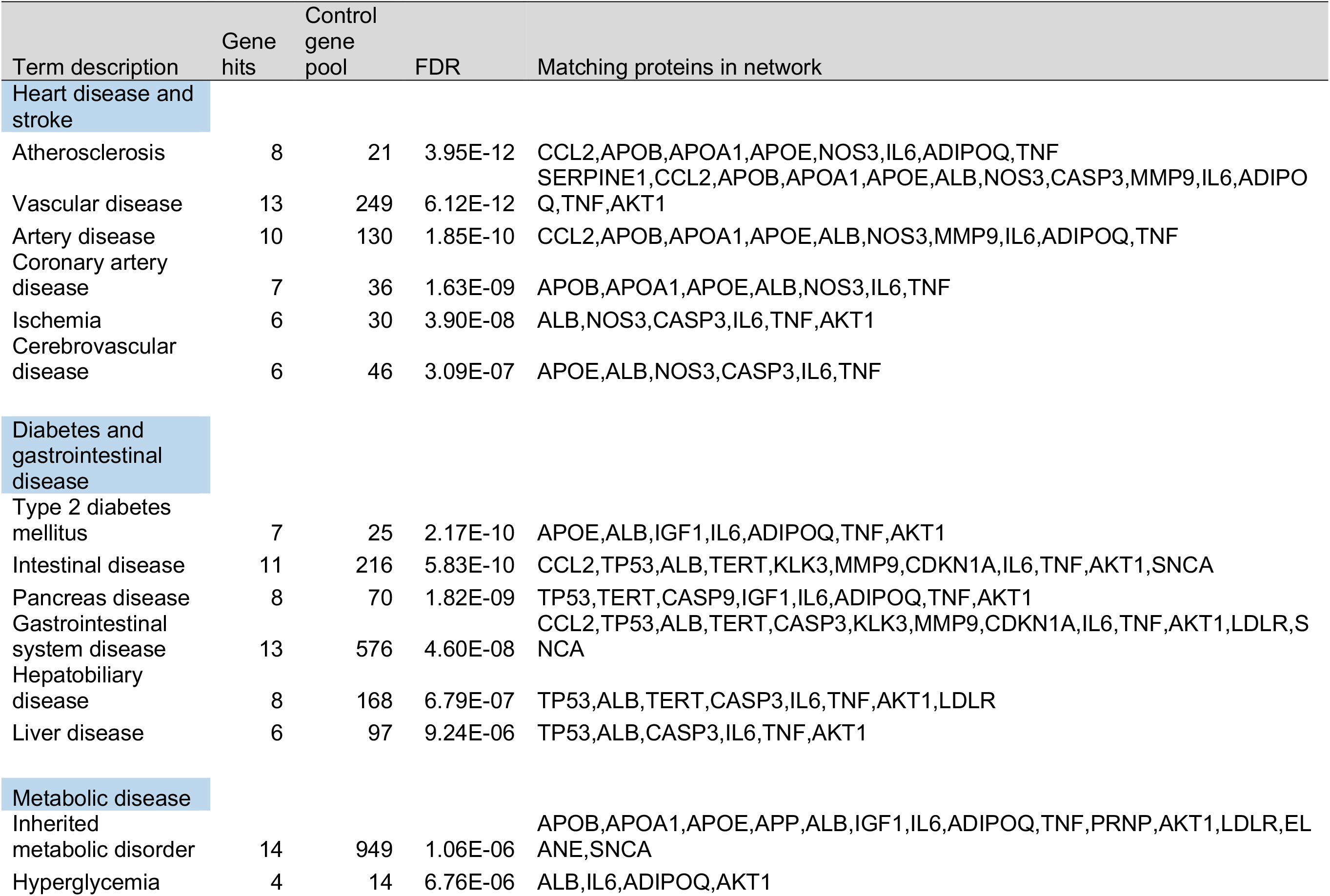
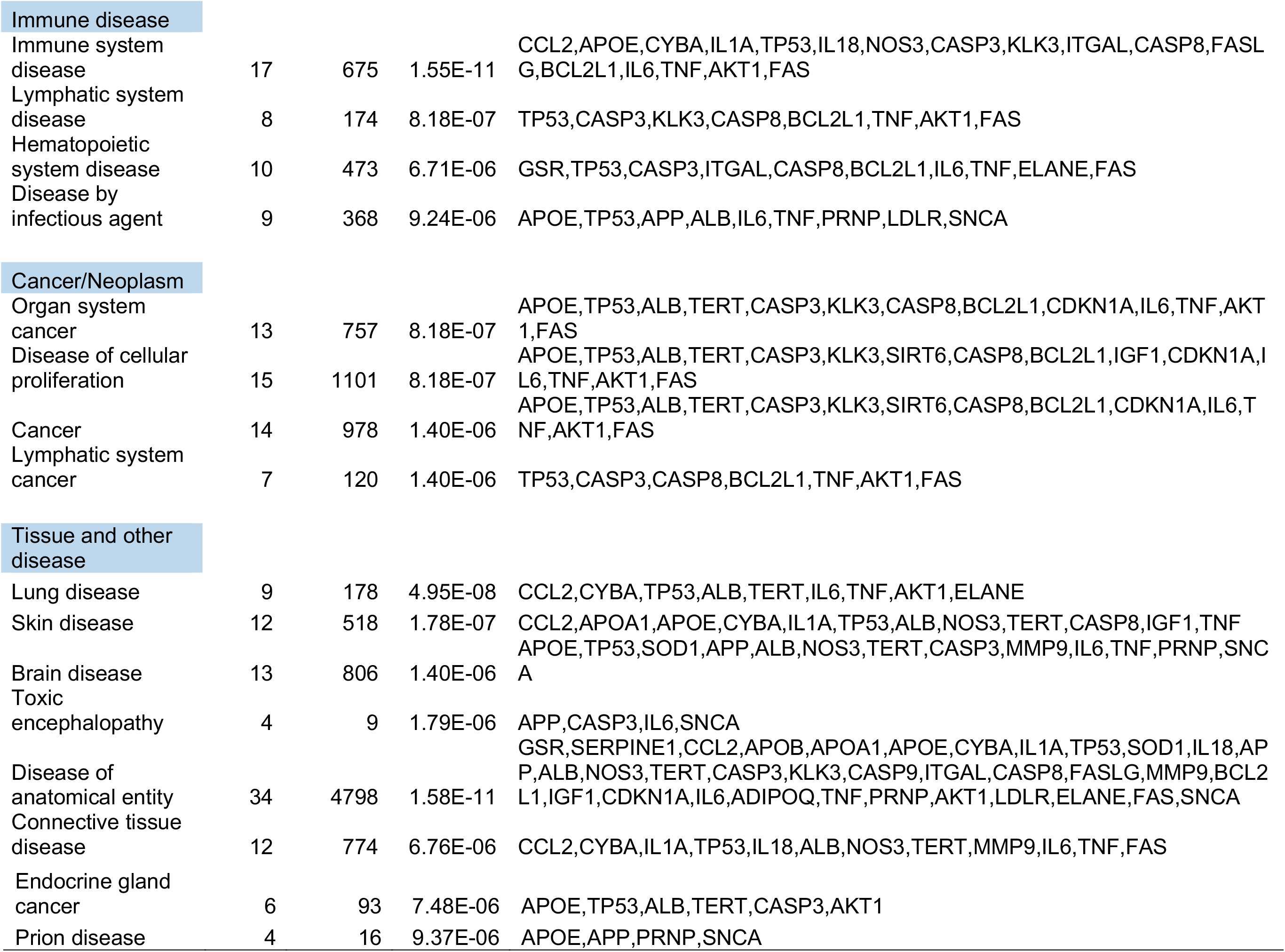
Disease association with the red-wine genes linked centenarians.

#### Tissue distribution

To elucidate the sites of action of the identified genes, we conducted a comprehensive analysis of the enriched tissue categories. Our findings revealed a significant enrichment of gene expression in specific tissues associated with the cardiovascular system (notably the blood and hematopoietic systems), gastrointestinal tract (particularly the liver and digestive tissues), and respiratory system (including lung and related respiratory tissues) (FDR < 3 × 10^−7^). These results underscore the concentration of gene activity within tissues integral to key physiological functions, such as digestion, circulation, and respiration, thereby supporting the biological pathways previously delineated (refer to the Discussion).

#### Diseases

Using disease association analysis, we identified 30 diseases exhibiting significant enrichment, with a false discovery rate (FDR) of less than 9.37 × 10^−6^ (Table 1). As illustrated in Table 1, the top ten diseases can be categorized as follows: (1) cardiovascular diseases, which represent heart disease and stroke, including atherosclerosis, vascular and arterial disease, and ischemic and cerebrovascular diseases, among others, with an FDR of less than 3.09 × 10^−7^; (2) type 2 diabetes, gastrointestinal diseases, and metabolic diseases, with an FDR of less than 9.24 × 10^−6^; (3) immune disease, with an FDR of less than 9.24 × 10^−6^; and (4) cancer, with an FDR of less than 1.04 × 10^−6^; and other diverse diseases.

## Discussion

Most human traits are shaped by intricate interactions among multiple genes, known as complex traits, rather than solely by individual genes (Fu et al., 2013; Buphamalai et al., 2021; Del Val et al., 2024). Gene network analysis is essential for mapping these complex interactions, significantly enhancing our understanding of their role in various phenotypic traits, including disease etiology. In this study, we identified specific genes linked to red wine and centenarians that underpin its longevity benefits. We present evidence of biological pathways, tissue distributions, and disease associations that highlight the advantages of red wine and its connection to exceptional longevity.

### Biological pathways

We used two methods to elucidate the hallmarks of centenarians and red wine. The ORA analysis showed general processes, mainly comprising response to stimuli, cell death/apoptosis, and metabolic processes (Figure 2B-C). The GSGE analysis provided an overlap but a more detailed biological ontology pathway enriched among the red wine genes with a stringent threshold of p-value < 1.00E-10. Upon reducing redundancies among these pathways, six major biological hallmark groups were delineated:

1. Regulation of smooth muscle cells
2. Regulation of neuronal apoptosis/death
3. Response to ultraviolet (UV) radiation (DNA damage response)
4. Response to lipopolysaccharide
5. Regulation of neuroinflammatory responses
6. Regulation of small-molecule metabolism (specifically lipoprotein metabolism)

The analysis of tissue distribution for these pathways revealed their essential roles across various biological systems, specifically highlighting three key areas: the gastrointestinal, cardiovascular, and respiratory systems. Each of these systems plays a crucial role in the relationship between red wine consumption and longevity. For example, in the gastrointestinal tract, the regulation of smooth muscle cells is vital for efficient food movement. In the cardiovascular system, these cells are important regulators of blood flow and blood pressure. Additionally, proper smooth muscle cell function is crucial for maintaining bronchiolar diameter in the respiratory system. This underscores the significance of smooth muscle regulation in various physiological contexts. Interestingly, there is an overlap between these tissues and the functions of the smooth muscle cells. Of the major sites of action (Hafen et al., 2025), three overlap with the areas (and functions of smooth muscle cells) of the gastrointestinal tract (propulsion of the food bolus), cardiovascular system (regulation of blood flow and pressure through vascular resistance), and respiratory tract (regulation of bronchiole diameter).

### Disease associations

This study also revealed significant connections with a wide range of 30 disease categories, highlighting a low false discovery rate (FDR) of less than 9.37 × 10^−6^. This level of precision suggests that the findings are robust.

The study categorizes diseases into six groups (Table 1) and finds that cardiovascular diseases, particularly heart disease and stroke, are significantly prevalent in the elderly population. Firstly, heart diseases refer to a range of health issues related to the heart and blood supply. These include atherosclerosis, vascular disease, arterial disease, coronary artery disease, and ischemia. These conditions are also linked to stroke in the brain and cerebrovascular disease (Brownstein et al., 2008; CDC, 2022). Notably, red wine together with Mediterranean diet and life-style interventions correlated with reduced risks of cognitive impairment, Alzheimer’s disease and Parkinson’s disease (Caruana et al., 2016; Nelson et al., 2024; Sakazaki et al., 2025), though red wine consumption as a single intervention does not seem effective (Reale et al., 2020).

Secondly, several categories can be combined, such as diabetes, gastrointestinal diseases, and metabolic disorders, particularly those affecting the pancreas and intestines. These three categories are closely interconnected (De Filippis et al., 2020; Adewuyi et al., 2024; Marathe et al., 2024). This suggests a potentially complex relationship in which red wine consumption may influence gut health and metabolic function, possibly due to the polyphenols found in red wine (Wang et al., 2021). Thirdly, the results highlight various immune disease categories, pointing to a wide range of health problems (Table 1). These include issues related to the hematopoietic, lymphatic, and immune systems, as well as blood cancers and infectious diseases (Janeway et al., 2001). Fourthly, the results identified a general cancer category that encompasses various types, specifically emphasizing organ system and lymphatic system cancers. Cancer is characterized as a disease marked by cellular proliferation. Finally, the findings revealed diverse tissue-specific diseases, ranging from endocrine disorders to conditions affecting peripheral tissues, such as the liver, skin, brain, and connective tissues, all linked to the cardiovascular system. Notably, our results also included prion diseases, which are associated with protein misfolding. The results indicate a strong association that may support the idea that red wine consumption positively influences cardiovascular and associated tissue health by influencing genetic pathways, which could play a protective role against these conditions.

### Methodological considerations

In this study, ORA and GSEA were used to analyze the genetic pathways related to red wine and centenarians. While the methods share the goal of identifying the biological significance of gene interactions, they differ in their approaches and the types of insights they provide. ORA focuses on determining whether a predefined set of genes exhibits statistical overrepresentation in a pool of genes of interest (Wieder et al., 2021; John et al., 2024). In contrast, GSEA takes a more comprehensive approach by analyzing the entire ranked list of genes rather than limiting the analysis to a specific subset (Mathur et al., 2018; Szklarczyk et al., 2025). Our results showed three biological ontology pathways in the ORA analysis (response to stimuli and cell death/apoptosis and metabolic processes), which is advantageous for examining more specific categories. The GSGA analysis showed broader groups, summarized into six pathway groups, which can be advantageous for an overview of the biological categories. This is consistent with the concept of the methodology, in which ORA provides a targeted examination of specific gene sets, whereas GSEA offers a broader view of gene expression dynamics across all genes, highlighting the interconnectedness of different pathways and processes.

#### Limitations

We conducted a comprehensive search that led us to identify the genes linked to both red wine consumption and the phenomenon of centenarians. However, upon closer examination, we discovered several discrepancies that highlighted the limitations of our study. Specifically, five of these genes (6.9 %) were solely associated with red wine consumption. Additionally, we identified 17 genes (23.6 %) that were exclusively associated with the characteristics of centenarians. Further complicating the results, there were seven genes (9.7 %) that showed no correlation with either red wine or centenarian status, leading to a total false-positive rate of 40.3%. We found that the AI chatbot, ChatGPT, showed 10% false positives, which was not accurate for validating the genes under the conditions used in this study.

These findings underscore the necessity for rigorous validation of the text search methodology, as the current results may not accurately reflect the true associations. Another notable limitation was the likelihood that our gene list was not fully saturated, suggesting that further research may be needed to uncover additional relevant genes and ensure a more comprehensive understanding of the relationship between red wine and longevity.

### Conclusions

This study explored the genetic factors linked to the health benefits of moderate red wine consumption, particularly its potential role in promoting longevity and healthy aging. It identified the genes associated with centenarians that may protect against a wide range of age-related diseases, including cardiovascular disease, diabetes, gastrointestinal diseases, metabolic disease, and cancer, among others. This analysis emphasizes the importance of these genes in biological processes, such as stress response and tissue health. While further research is needed to deepen the understanding of these genetic influences, the findings suggest a genetic basis for the benefits of red wine, where moderate consumption contributes to a healthier lifespan. The discussion highlights the need for a more comprehensive and multidisciplinary approach to aging, focusing on the biological characteristics of centenarians and the effects of red wine to identify potential therapeutic strategies. Overall, the study highlights intriguing associations that merit further investigation, especially to understand the biological mechanisms at play and whether these associations can inform dietary recommendations or therapeutic approaches in preventing and bypassing diverse diseases.

## Acknowledgments

We would like to extend our gratitude to the members of the Murakami Laboratory for their invaluable technical assistance and helpful discussions. The manuscript has been refined for clarity and readability through the collaborative efforts of the team, as well as the use of grammar-checking software “https://app.grammarly.com (last accessed on 20 June 2025).”

## Notes

### Competing Interest Statement

The authors have declared no competing interest.

## References

1. Adewuyi EO, Porter T, O’Brien EK, Olaniru O, Verdile G, Laws SM. Genome-wide cross-disease analyses highlight causality and shared biological pathways of type 2 diabetes with gastrointestinal disorders. Commun Biol. 2024;7(1):643. Published 2024 May 27. doi:10.1038/s42003-024-06333-z

2. Badial K, Lacayo P, Murakami S. Biology of Healthy Aging: Biological Hallmarks of Stress Resistance Related and Unrelated to Longevity in Humans. Int J Mol Sci. 2024;25(19):10493. Published 2024 Sep 29. doi:10.3390/ijms251910493

3. Balmorez T, Sakazaki A, Murakami S. Genetic Networks of Alzheimer’s Disease, Aging, and Longevity in Humans. Int J Mol Sci. 2023;24(6):5178. Published 2023 Mar 8. doi:10.3390/ijms24065178

4. Bernatoniene J, Kopustinskiene DM. The Role of Catechins in Cellular Responses to Oxidative Stress. Molecules. 2018 Apr 20;23(4):965. doi: 10.3390/molecules23040965. PMID: 29677167; PMCID: PMC6017297.

5. Bonnefont-Rousselot, D. (2016). Resveratrol and Cardiovascular Diseases. Nutrients, 8(5), 250. 10.3390/nu8050250

6. Brownstein JN. Addressing heart disease and stroke prevention through comprehensive population-level approaches. Prev Chronic Dis. 2008;5(2):A31.

7. Brunet A. Old and new models for the study of human ageing. Nat Rev Mol Cell Biol. 2020 Sep;21(9):491–493. doi: 10.1038/s41580-020-0266-4. PMID: 32572179; PMCID: PMC7531489.

8. Buphamalai P, Kokotovic T, Nagy V, Menche J. Network analysis reveals rare disease signatures across multiple levels of biological organization. Nat Commun. 2021;12(1):6306. Published 2021 Nov 9. doi:10.1038/s41467-021-26674-1

9. Caruana M, Cauchi R, Vassallo N. Putative Role of Red Wine Polyphenols against Brain Pathology in Alzheimer’s and Parkinson’s Disease. Front Nutr. 2016;3:31. Published 2016 Aug 12. doi:10.3389/fnut.2016.00031

10. Centers for Disease Control and Prevention. Best Practices for Heart Disease and Stroke: A Guide to Effective Approaches and Strategies; 2022. doi:10.15620/cdc:122290.

11. De Filippis A, Ullah H, Baldi A, et al. Gastrointestinal Disorders and Metabolic Syndrome: Dysbiosis as a Key Link and Common Bioactive Dietary Components Useful for their Treatment. Int J Mol Sci. 2020;21(14):4929. Published 2020 Jul 13. doi:10.3390/ijms21144929

12. Del Val C, Díaz de la Guardia-Bolívar E, Zwir I, et al. Gene expression networks regulated by human personality. Mol Psychiatry. 2024;29(7):2241–2260. doi:10.1038/s41380-024-02484-x

13. Dong Y, Duan S, Xia Q, Liang Z, Dong X, Margaryan K, Musayev M, Goryslavets S, Zdunić G, Bert PF, Lacombe T, Maul E, Nick P, Bitskinashvili K, Bisztray GD, Drori E, De Lorenzis G, Cunha J, Popescu CF, Arroyo-Garcia R, Arnold C, Ergül A, Zhu Y, Ma C, Wang S, Liu S, Tang L, Wang C, Li D, Pan Y, Li J, Yang L, Li X, Xiang G, Yang Z, Chen B, Dai Z, Wang Y, Arakelyan A, Kuliyev V, Spotar G, Girollet N, Delrot S, Ollat N, This P, Marchal C, Sarah G, Laucou V, Bacilieri R, Röckel F, Guan P, Jung A, Riemann M, Ujmajuridze L, Zakalashvili T, Maghradze D, Höhn M, Jahnke G, Kiss E, Deák T, Rahimi O, Hübner S, Grassi F, Mercati F, Sunseri F, Eiras-Dias J, Dumitru AM, Carrasco D, Rodriguez-Izquierdo A, Muñoz G, Uysal T, Özer C, Kazan K, Xu M, Wang Y, Zhu S, Lu J, Zhao M, Wang L, Jiu S, Zhang Y, Sun L, Yang H, Weiss E, Wang S, Zhu Y, Li S, Sheng J, Chen W. Dual domestications and origin of traits in grapevine evolution. Science. 2023 Mar 3;379(6635):892–901. doi: 10.1126/science.add8655. Epub 2023 Mar 2. PMID: 36862793.

14. Elizarraras JM, Liao Y, Shi Z, Zhu Q, Pico AR, Zhang B. WebGestalt 2024: faster gene set analysis and new support for metabolomics and multi-omics. Nucleic Acids Res. 2024;52(W1):W415–W421. doi:10.1093/nar/gkae456

15. Ferrières J. The French paradox: lessons for other countries. Heart. 2004 Jan;90(1):107–11. doi: 10.1136/heart.90.1.107. PMID: 14676260; PMCID: PMC1768013.

16. Fischetti M, Franchi F. Wine’s True Origins: A broad genetic study has revised the prevailing narrative about how wine grapes spread around the world. Sci Am. 2023 Oct 1;329(3):38. doi: 10.1038/scientificamerican1023-38. PMID: 39017261.

17. Fu W, O’Connor TD, Akey JM. Genetic architecture of quantitative traits and complex diseases. Curr Opin Genet Dev. 2013 Dec;23(6):678–83. doi: 10.1016/j.gde.2013.10.008. Epub 2013 Nov 26. PMID: 24287334; PMCID: PMC6764439.

18. Hafen BB, Shook M, Burns B. Anatomy, Smooth Muscle. [Updated 2023 Jul 17]. In: StatPearls [Internet]. Treasure Island (FL): StatPearls Publishing; 2025 Jan-. Available from: https://www.ncbi.nlm.nih.gov/books/NBK532857/

19. Haseeb S, Alexander B, Santi RL, Liprandi AS, Baranchuk A. What’s in wine? A clinician’s perspective. Trends Cardiovasc Med. 2019 Feb;29(2):97–106. doi: 10.1016/j.tcm.2018.06.010. Epub 2018 Jun 26. PMID: 30104174.

20. Hrelia S, Di Renzo L, Bavaresco L, Bernardi E, Malaguti M, Giacosa A. Moderate Wine Consumption and Health: A Narrative Review. Nutrients. 2022 Dec 30;15(1):175. doi: 10.3390/nu15010175. PMID: 36615832; PMCID: PMC9824172.

21. Janeway CA Jr, Travers P, Walport M, et al. Immunobiology: The Immune System in Health and Disease. 5th edition. New York: Garland Science; 2001. The components of the immune system. Available from: https://www.ncbi.nlm.nih.gov/books/NBK27092/

22. John M Elizarraras, Yuxing Liao, Zhiao Shi, Qian Zhu, Alexander R Pico, Bing Zhang, WebGestalt 2024: faster gene set analysis and new support for metabolomics and multi-omics, Nucleic Acids Research, 2024, gkae456

23. Lucerón-Lucas-Torres M, Cavero-Redondo I, Martínez-Vizcaíno V, Bizzozero-Peroni B, Pascual-Morena C, Álvarez-Bueno C. Association between wine consumption and cancer: a systematic review and meta-analysis. Front Nutr. 2023 Sep 4;10:1197745. doi: 10.3389/fnut.2023.1197745. PMID: 37731399; PMCID: PMC10507274.

24. Lucerón-Lucas-Torres M, Cavero-Redondo I, Martínez-Vizcaíno V, Saz-Lara A, Pascual-Morena C, Álvarez-Bueno C. Association Between Wine Consumption and Cognitive Decline in Older People: A Systematic Review and Meta-Analysis of Longitudinal Studies. Front Nutr. 2022 May 12;9:863059. doi: 10.3389/fnut.2022.863059. PMID: 35634389; PMCID: PMC9133879.

25. Lucerón-Lucas-Torres M, Saz-Lara A, Díez-Fernández A, Martínez-García I, Martínez-Vizcaíno V, Cavero-Redondo I, Álvarez-Bueno C. Association between Wine Consumption with Cardiovascular Disease and Cardiovascular Mortality: A Systematic Review and Meta-Analysis. Nutrients. 2023 Jun 17;15(12):2785. doi: 10.3390/nu15122785. PMID: 37375690; PMCID: PMC10303697.

26. Machino K, Link CD, Wang S, Murakami H, Murakami S. A semi-automated motiontracking analysis of locomotion speed in the C. elegans transgenics overexpressing beta-amyloid in neurons. Front Genet. 2014;5:202. Published 2014 Jul 4. doi:10.3389/fgene.2014.00202

27. Marathe CS, Rayner CK, Wu T, et al. Gastrointestinal Disorders in Diabetes. [Updated 2024 Feb 22]. In: Feingold KR, Ahmed SF, Anawalt B, et al., editors. Endotext [Internet]. South Dartmouth (MA): MDText.com, Inc.; 2000-. Available from: https://www.ncbi.nlm.nih.gov/books/NBK553219/

28. Markoski MM, Garavaglia J, Oliveira A, Olivaes J, Marcadenti A. Molecular Properties of Red Wine Compounds and Cardiometabolic Benefits. Nutr Metab Insights. 2016 Aug 2;9:51–7. doi: 10.4137/NMI.S32909. PMID: 27512338; PMCID: PMC4973766.

29. Mathur R, Rotroff D, Ma J, Shojaie A, Motsinger-Reif A. Gene set analysis methods: a systematic comparison. BioData Min. 2018;11:8. Published 2018 May 31. doi:10.1186/s13040-018-0166-8

30. Murakami S, Lacayo P. Biological and disease hallmarks of Alzheimer’s disease defined by Alzheimer’s disease genes. Front Aging Neurosci. 2022;14:996030. Published 2022 Nov 9. doi:10.3389/fnagi.2022.996030

31. Nielson KA, Venneri A, Murakami S. Editorial: Insights in neurocognitive aging and behavior: 2022. Front Aging Neurosci. 2024;16:1361839. Published 2024 Jan 16. doi:10.3389/fnagi.2024.1361839

32. Reale M, Costantini E, Jagarlapoodi S, Khan H, Belwal T, Cichelli A. Relationship of Wine Consumption with Alzheimer’s Disease. Nutrients. 2020;12(1):206. Published 2020 Jan 13. doi:10.3390/nu12010206

33. Robin-Champigneul F. Jeanne Calment’s Unique 122-Year Life Span: Facts and Factors; Longevity History in Her Genealogical Tree. Rejuvenation Res. 2020;23(1):19–47. doi:10.1089/rej.2019.2298

34. Robine JM, Allard M, Herrmann FR, Jeune B. The Real Facts Supporting Jeanne Calment as the Oldest Ever Human. J Gerontol A Biol Sci Med Sci. 2019;74(Suppl_1):S13–S20. doi:10.1093/gerona/glz198

35. Rocca C, Angelone T. Editorial commentary: A chemically complex and unique beverage: The wine. Trends Cardiovasc Med. 2019 Feb;29(2):107–108. doi: 10.1016/j.tcm.2018.07.008. Epub 2018 Jul 19. PMID: 30075907.

36. Sakazaki A, Lui A, Muhammad H, Vu C, le Roux A, Lacayo P, Murakami S. Minimum Physical Activities Protective Against Alzheimer’s Disease in Late Life: A Systematic Review. medRxiv 2025.04.01.25325006; doi: 10.1101/2025.04.01.25325006

37. Snopek L, Mlcek J, Sochorova L, Baron M, Hlavacova I, Jurikova T, Kizek R, Sedlackova E, Sochor J. Contribution of Red Wine Consumption to Human Health Protection. Molecules. 2018 Jul 11;23(7):1684. doi: 10.3390/molecules23071684. PMID: 29997312; PMCID: PMC6099584.

38. Sziraki A, Lu Z, Lee J, Banyai G, Anderson S, Abdulraouf A, Metzner E, Liao A, Banfelder J, Epstein A, Schaefer C, Xu Z, Zhang Z, Gan L, Nelson PT, Zhou W, Cao J. A global view of aging and Alzheimer’s pathogenesis-associated cell population dynamics and molecular signatures in human and mouse brains. Nat Genet. 2023 Dec;55(12):2104–2116. doi: 10.1038/s41588-023-01572-y. Epub 2023 Nov 30. PMID: 38036784; PMCID: PMC10703679.

39. Szklarczyk D, Nastou K, Koutrouli M, et al. The STRING database in 2025: protein networks with directionality of regulation. Nucleic Acids Res. 2025;53(D1):D730–D737. doi:10.1093/nar/gkae1113

40. Tenopir, C. (2008). Online Systems for Information Access and Retrieval. Library Trends 56(4), 816-829. 10.1353/lib.0.0005.

41. Vahdati Nia B, Kang C, Tran MG, Lee D, Murakami S. Meta Analysis of Human AlzGene Database: Benefits and Limitations of Using C. elegans for the Study of Alzheimer’s Disease and Co-morbid Conditions. Front Genet. 2017;8:55. Published 2017 May 12. doi:10.3389/fgene.2017.00055

42. Wang K, Sun G, Conlon MA, Ren W, Yang G. Editorial: Dietary Polyphenols for Improving Gut Health: Volume 1. Front Nutr. 2021;8:760917. Published 2021 Oct 4. doi:10.3389/fnut.2021.760917

43. Wieder C, Frainay C, Poupin N, Rodríguez-Mier P, Vinson F, Cooke J, Lai RP, Bundy JG, Jourdan F, Ebbels T. Pathway analysis in metabolomics: Recommendations for the use of over-representation analysis. PLoS Comput Biol. 2021 Sep 7;17(9):e1009105. doi: 10.1371/journal.pcbi.1009105. PMID: 34492007; PMCID: PMC8448349.

44. Wu QJ, Zhang TN, Chen HH, et al. The sirtuin family in health and disease. Signal Transduct Target Ther. 2022;7(1):402. Published 2022 Dec 29. doi:10.1038/s41392-022-01257-8

